# The anterior insula channels prefrontal expectancy signals during affective processing

**DOI:** 10.1101/536300

**Authors:** Vanessa Teckentrup, Johan N. van der Meer, Viola Borchardt, Yan Fan, Monja P. Neuser, Claus Tempelmann, Luisa Herrmann, Martin Walter, Nils B. Kroemer

**Affiliations:** University of Tübingen, Department of Psychiatry and Psychotherapy, Tübingen, Germany; Queensland Institute of Medical Research, Brisbane, Australia; University of Magdeburg, Department of Psychiatry and Psychotherapy, Germany; Clinical Affective Neuroimaging Laboratory, Magdeburg, Germany; Leibniz Institute for Neurobiology, Magdeburg, Germany; Leibniz Research Centre for Working Environment and Human Factors, Department of Psychology and Neurosciences Dortmund, German; University of Magdeburg, Department of Neurology, Germany

**Author notes:** Corresponding authors: Prof. Dr. Martin Walt, Dr. Nils B. Kroemer, Department of Psychiatry and Psychotherapy, University of Tübingen, Calwerstr. 14, 72076 Tübingen, Germany. contributed equally (Martin Walter, Nils B. Kroemer).

**Keywords:** signaling dynamics, cross-correlation, functional networks, fMRI, expectancy, hierarchical linear modeling

## Abstract

Expectancy shapes our perception of impending events. Although such an interplay between cognitive and affective processes is often impaired in mental disorders, it is not well understood how top-down expectancy signals modulate future affect. We therefore track the information flow in the brain during cognitive and affective processing segregated in time using task-specific cross-correlations. Participants in two independent fMRI studies (N_1_ = 37 & N_2_ = 55) were instructed to imagine a situation with affective content as indicated by a cue, which was then followed by an emotional picture congruent with expectancy. To correct for intrinsic covariance of brain function, we calculate resting-state cross-correlations analogous to the task. First, using factorial modeling of delta cross-correlations (task-rest) of the first study, we find that the magnitude of expectancy signals in the anterior insula cortex (AIC) modulates the BOLD response to emotional pictures in the anterior cingulate and dorsomedial prefrontal cortex in opposite directions. Second, using hierarchical linear modeling of lagged connectivity, we demonstrate that expectancy signals in the AIC indeed foreshadow this opposing pattern in the prefrontal cortex. Third, we replicate the results in the second study using a higher temporal resolution, showing that our task-specific cross-correlation approach robustly uncovers the dynamics of information flow. We conclude that the AIC arbitrates the recruitment of distinct prefrontal networks during cued picture processing according to triggered expectations. Taken together, our study provides new insights into neuronal pathways channeling cognition and affect within well-defined brain networks. Better understanding of such dynamics could lead to new applications tracking aberrant information processing in mental disorders.

## Introduction

Whenever we are confronted with a stirring situation, our prior beliefs and expectations will influence future perception. For example, if you have to give a talk, the preparation can be associated with different expectations on how it will resonate with the audience. In turn, this will shape your perception of the listeners: If you are affected by stage fright, your expectations are likely negative and you may interpret an ambiguous facial expression as disapproving. In contrast, if you expect the audience to respond well, you may interpret the same expression as showing focused attention. Although such processes often come into play in our daily life (de Lange, Heilbron, & Kok, 2018) and aberrant expectations play an important role in affective disorders (Disner, Beevers, Haigh, & Beck, 2011; Holtzheimer & Mayberg, 2011), the exact neural processes subserving the channeling of expectancy are not well understood to date.

So far, neuroimaging research on cognitive-affective processing has largely focused on either delineating neural circuits encoding cognitive and affective processes or studying the affective modulation of cognitive processing. On the one hand, brain regions involved in either expectancy or emotional-picture processing have been identified. For example, pioneering studies (Bermpohl et al., 2006b) have suggested that expectancy primarily recruits regions in the parieto-occipital sulcus and anterior cingulate. In contrast, affective processing led to increased activation in prefrontal as well as limbic regions. On the other hand, brain regions associated with the simultaneous representation of attention and affect have been identified (Dolan, 2002; J. R. Gray, Braver, & Raichle, 2002; Ho, Gonzalez, Abelson, & Liberzon, 2012; Pessoa, 2008, 2016) pointing to the dorsomedial prefrontal cortex (dmPFC) as a prime candidate for the arbitration of expectancy (Bermpohl et al., 2006a; Walter et al., 2009). The dmPFC is involved in a variety of related task domains ranging from emotional judgement (Northoff et al., 2004) over memory retrieval (Macrae, Moran, Heatherton, Banfield, & Kelley, 2004) to decision making (Ruff & Fehr, 2014), suggesting a role as hub in processing of task demands. In line with this, hyperconnectivity of the dmPFC with other key networks has been shown in patients suffering from major depressive disorder (MDD) compared to healthy controls (Sheline, Price, Yan, & Mintun, 2010). Congruously, marked differences in dmPFC signaling during expected versus unexpected picture viewing between MDD patients and healthy controls were previously shown (Bermpohl et al., 2009; Zhang et al., 2017), illustrating aberrant interplay between cognitive and affective processes in depression. Thus, whereas the mediating role of the dmPFC has been widely established, it is still an open question how prior information encoded during the cue phase is carried over to inform picture processing as the link between both phases has not been detailed so far.

In the past decade, studies moved from mapping functions to specific regions to well-defined networks instead, defined by correlated BOLD responses reflecting intrinsic functional connectivity. Task-negative (or internally-oriented) processes are primarily associated with the default mode network (DMN) comprising the ventromedial prefrontal and the posterior cingulate cortex as key nodes. In contrast, task-positive (or externally-oriented) processes are primarily associated with the central executive network (CEN) comprising dorsomedial and dorsolateral prefrontal cortices, the frontal eye fields as well as the posterior parietal cortex as key nodes (Bressler & Menon, 2010). The salience network (SN) orchestrates the switching between internally versus externally oriented processes and comprises the dorsal anterior cingulate cortex and the AIC as key nodes. In line with its key role in arbitrating task-positive versus task-negative components, differences in the SN have been related to various dysfunctions in mental disorders (Menon, 2011; Uddin, 2015). Critically, separate cue and picture processing phases help to differentiate task-negative versus task-positive brain networks as processing of expectancy cues is primarily based on (internal) mentalizing whereas picture processing relies on external affective input. Such a complementary network architecture is further substantiated in Embodied Predictive Interoceptive Coding (EPIC model, Feldman Barrett & Simmons, 2015). According to the EPIC model, regions within the DMN generate expectations of upcoming stimuli based on previous experiences whereas regions within the CEN (i.e., the dmPFC) compute prediction errors in matching sensations with expectations. Hence, task-negative (i.e. interoceptive) networks may form expectations based on cues that are evaluated by task-positive, prediction error networks, but how task-specific dynamics unfold within these functional networks remains to be tested.

Consequently, the idea to capture temporal dynamics in neural information processing has gained considerable traction recently (Buckner, Krienen, & Yeo, 2013; Hutchison, Womelsdorf, Gati, Everling, & Menon, 2013; Preti, Bolton, & Van De Ville, 2017). Whereas dynamic functional connectivity analyses of resting-state data has shown the emergence of coordinated functional networks even in the absence of a dedicated task (Deco, Jirsa, & McIntosh, 2011; Hutchison et al., 2013), these studies are necessarily limited in uncovering causes for shifts in connectivity due to the absence of experimental control over events (Gonzalez-Castillo & Bandettini, 2017; Krakauer, Ghazanfar, Gomez-Marin, MacIver, & Poeppel, 2017). In contrast, a task provides temporal structure allowing for more nuanced testing of the link between cognition and affect. Across time such a link can be studied via cross-correlation analysis. To calculate cross-correlation, a set of time-shifted (lagged) versions of a time series is correlated with a second (unshifted) time series (*Figure 1*). This technique has been widely used to study the neuronal signal transduction in single- or multi-unit recordings of the cat visual cortex (C. M. Gray, Konig, Engel, & Singer, 1989; Innocenti, Lehmann, & Houzel, 1994; Munk, Nowak, Nelson, & Bullier, 1995; Nowak, Munk, Nelson, James, & Bullier, 1995; Schwarz & Bolz, 1991). Crucially, although cross-correlations reliably uncovered the temporal structure of signal transduction (Munk et al., 1995; Nowak et al., 1995), its use remained primarily restricted to coupling between single neurons. To explore the functional coupling of networks over time as measured with fMRI, we employed cross-correlation analysis to track information flow from regions encoding expectancy to prefrontal networks encoding affective pictures.

**Figure 1.**
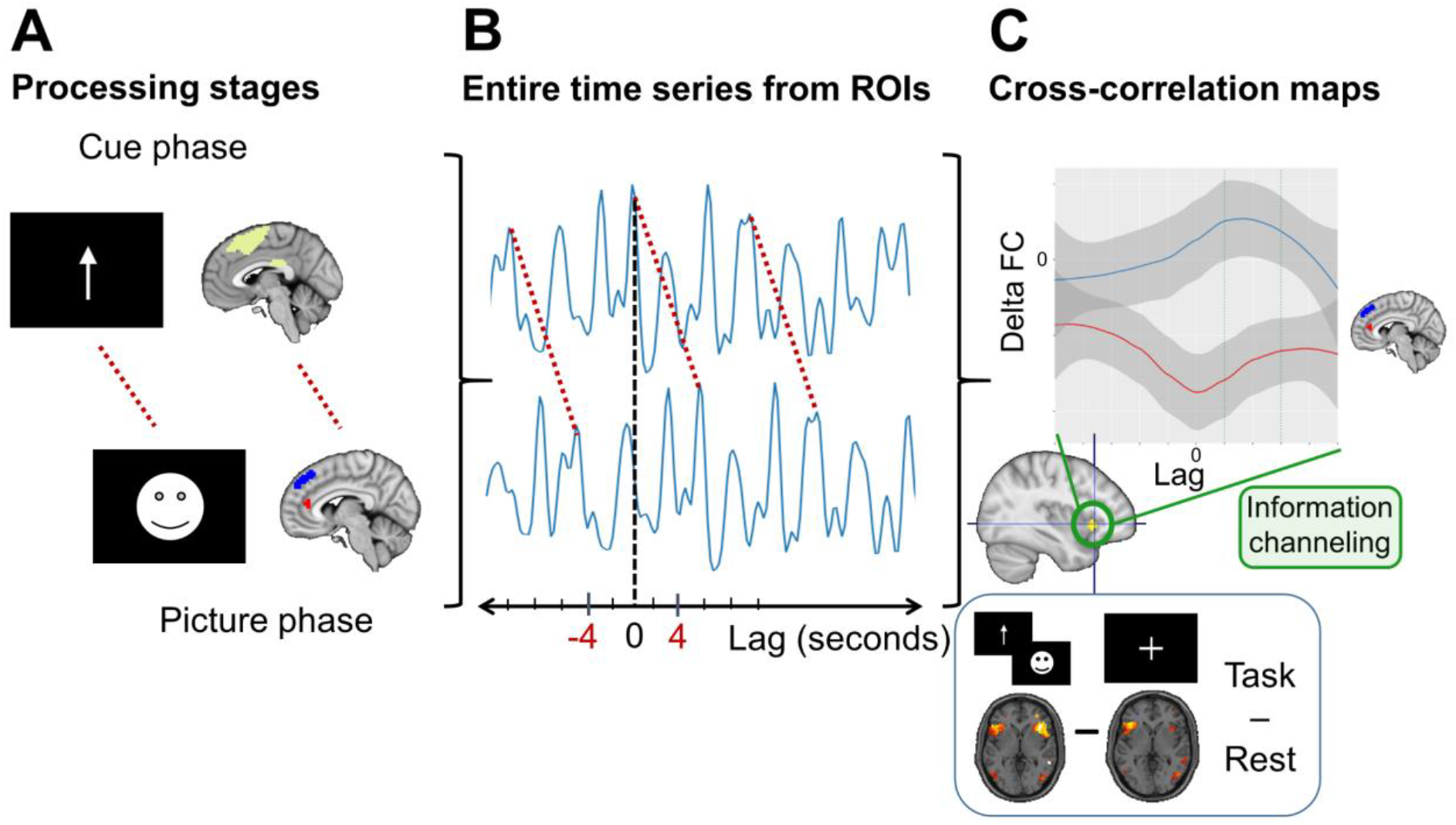
Data-analysis workflow with (A) definition of the cue phase related mask and the ROIs associated with the picture processing phase. Due to bioRxiv’s policy to avoid the inclusion of photographs and any other identifying information of people, stimulus pictures were replaced for this figure. (B) Time series from the mask (voxel-wise) and ROIs (mean time-series) associated with task-positive (dmPFC, blue) and task-negative (pgACC, red) networks were extracted and cross-correlated. Cross-correlation maps were generated for task and resting-state data in the same way. (C) Resting-state cross-correlation maps were subtracted from task-based cross-correlation maps to create time-lagged delta functional connectivity (dFC) maps which were submitted to statistical analyses. We hypothesized the anterior insula (AIC) to act as a switch, modulating BOLD responses in task-positive and task-negative regions. This channeling of information based on the preceding cue would be reflected by a modulatory connectivity profile over time between the AIC and the two picture phase ROIs (dmPFC and pgACC), respectively.

To summarize, although extensive work has recently uncovered the dynamic interplay between functional networks at rest, less is known about task-inherent event-related dynamics that are key in affective processing. To bridge this gap, we employ cross-correlation analysis to determine the temporal dynamics in the brain reflecting cue-induced expectations and their neural traces during emotional processing. Thus, we expected BOLD signals in areas involved in cue processing to foreshadow expectancy traces in prefrontal regions associated with cognitive-emotional integration. Accordingly, in two independent datasets, our cross-correlation approach identified the AIC as the key switch channeling BOLD response to emotional pictures towards either DMN (anterior cingulate) or CEN (dmPFC) nodes indicating opposing processing streams in the prefrontal cortex.

## Methods

### Participants and procedure

We conducted two independent functional MRI studies. In the first study (exploration: EXP), 37 healthy males were included in the current analysis (M_age_ = 43.7 years ± 9.8, range: 31-59). Two participants had to be excluded due to excessive motion. Participants completed the emotional expectancy task and resting-state sessions as part of a clinical trial (NCT02602275) where after baseline measurements, subjects took either placebo or verum (Neurexan) in a counterbalanced order. Neurexan (Nx4) is a medicinal product, consisting of three herbal extracts (Avena sativa, Coffea arabica, Passiflora incarnate) and one mineral salt (Zincum isovalericum). Data was recorded with a simultaneous EEG-fMRI setup in which EEG was recorded during fMRI scans using a BrainAmp MR system (Brain Products) with a 64-channel Easycap augmented with six carbon-wire loops (CWLs) (Sikka et al., submitted; van der Meer et al., 2016). Due to session effects when participants complete the task for a second time, we restricted our analyses to the first day and pooled the data across conditions (verum vs. placebo) to establish the method orthogonal to treatment.

In Study two (replication: REP), we included 55 healthy participants (13 female, M_age_ = 32.1 years ± 8.5, range: 22-52). One participant had to be excluded due to excessive motion. Participants in this study completed a resting-state fMRI measurement (10 minutes) before the salience expectancy task (14 minutes; for details, see SI).

### Paradigm

In our exploration study, we employed an emotional expectancy task adapted from Walter et al. (2009) which included passive cue and picture viewing (*Figure 1A*). The paradigm was based on a factorial design with the two factors *expectancy* (expected, unexpected) and *emotion* (positive, negative, and neutral). Every trial started with a fixation cross which was shown for a jittered interval between 4 and 6.5 seconds. In half of the trials, the fixation cross was followed by the cue phase, in which the valence of the subsequently presented picture was cued by an arrow (positive: upwards pointing, negative: downwards pointing, neutral: pointing to the right) for a varying duration of 3-5 seconds. To generate an expectation of a following stimulus presentation, participants were instructed to envision a situation congruent with the cue. In the subsequent picture phase, a picture congruent to the cued valence was shown for 4 seconds. In the other half of the trials, unexpected pictures were not preceded by a cue phase but were shown directly after the fixation cross. In total, 60 pictures from the International Affective Picture System (Lang, Bradley, & Cuthbert, 2008) were used in a counterbalanced design (i.e., 20 pictures per valence category). Completing the paradigm took about 15 minutes.

In the replication study, we used a similar paradigm described in detail elsewhere (Li et al., 2017). Briefly, the design was modified to show expected or unexpected emotional pictures of low or high salience. Moreover, an exclamation mark appeared next to the arrow as well as points indicating how many objects will be shown on screen (for details, see SI).

### MRI data acquisition and preprocessing

For the exploration study, structural and functional MRI data was acquired on a 3T Philips Achieva magnetic resonance imaging scanner. Structural T1-weighted images were measured using a Turbo Field Echo sequence with 274 sagittal slices covering the whole brain, flip angle = 8°, 256 × 256 matrix size and voxel size = 0.7 × 0.7 × 0.7 mm^3^. The functional MRI data were T2*-weighted echo-planar images (EPIs) with 429 volumes, 34 axial slices covering the whole brain, repetition time (TR) = 2 s, echo time (TE) = 30 ms, flip angle = 90°, 96 × 94 matrix, field of view = 240 × 240 mm^2^ and voxel size = 2.5 × 2.5 × 3 mm^3^. The resting-state data consisted of 355 volumes of T2*-weighted echo-planar images (EPIs) that were measured with the same sequence parameters as the task data.

For the replication study, structural and functional MRI data was acquired on a 3T Siemens Trio magnetic resonance imaging scanner. Structural T1-weighted images were measured using a MPRAGE sequence with 192 sagittal slices covering the whole brain, flip angle = 7°, 256 × 256 matrix size and voxel size = 1 × 1 × 1 mm^3^. As the main interest of this study was the temporal decomposition of dynamic network changes, the functional MRI data for the replication was acquired with a higher temporal, but lower spatial resolution. Thus, the functional data comprised 672 volumes of T2*-weighted echo-planar images (EPIs), 26 axial slices covering the whole brain, TR = 1.25 s, TE = 25 ms, flip angle = 70°, 44 × 44 matrix, field of view = 220 × 220 mm^2^ and voxel size = 5 × 5 × 5 mm^3^. The resting-state data consisted of 478 volumes of T2*-weighted EPIs that were measured with the same sequence parameters as the task data.

### MRI data preprocessing

Task fMRI data was submitted to SPM12 (Statistical parametric mapping, Wellcome Department of Imaging Neuroscience, London, UK) using MATLAB (The Mathworks Inc., Natick, MA, USA). First, slice timing was corrected for each volume by interpolating the slices to the middle slice, then all volumes were realigned to the first volume by applying a rigid-body transformation to correct for head motion. Participants with head movement exceeding 3 mm for translation head motion parameters or 3° for rotation head motion parameters were excluded from further analyses. The anatomical images were co-registered to match the functional images. All images were then spatially normalized to standard MNI space following tissue segmentation of the anatomical images and subsequent application of the generated deformation field to the functional images. All images were smoothed with a Gaussian kernel with 8 mm FWHM.

Resting-state (rs-) fMRI data was preprocessed using SPM12 and DPABI (http://rfmri.org/dpabi). The first 5 volumes were discarded to allow the MR signal to achieve T1 equilibrium. Then, slice timing was corrected for each volume by interpolating the slices to the middle slice and all volumes were realigned to the first volume by applying a rigid-body transformation to correct for head motion. The anatomical images were co-registered to match the functional images, then segmented into gray matter and white matter. Subject-specific templates were created with diffeomorphic anatomical registration using DARTEL (Ashburner, 2007). A group-specific template was then created from all subject-specific templates. Co-registered rs-fMRI data were subsequently normalized to the MNI template. Physiological noise was reduced by regressing out signals from white matter, cerebrospinal fluid and the 6-rigid body realignment parameters. All images were smoothed with a Gaussian kernel with 8 mm FWHM. Importantly, global signal removal was not performed to avoid false induction of anti-correlations between time-series (Murphy, Birn, Handwerker, Jones, & Bandettini, 2009). For more detailed information, see SI.

### Definition of regions of interest (ROI)

To capture the representation of general expectancy (collapsed over valence) during the cue phase, a mask was defined that represented voxels showing activation within the cue phase compared to baseline. Based on the linear association of cue representation with the contrast expected picture viewing > unexpected picture viewing, we defined a cluster in the pregenual anterior cingulate cortex (pgACC) that represented the task-negative (or internally-driven) aspect of affective picture processing. We also hypothesized that the dmPFC plays a key role in handling affective picture processing given its continuous recruitment in simultaneous processing of cognitive and emotional demands (Bermpohl et al., 2006a; Walter et al., 2009). Thus, we selected a second ROI in the dmPFC taken from Walter et al. (2009). To ensure comparable analysis pathways for the exploration and the replication datasets, all masks were resliced to the dimensions and voxel size of the replication datasets. A detailed description of mask and ROI generation can be found in the SI.

### Time-series data extraction

For the subsequent calculation of the time-shifted functional connectivities (FC, *Figure 1B*), the time-series of each voxel inside the three different masks (cue phase mask, picture phase / pgACC mask and picture phase / dmPFC mask) was extracted from the preprocessed resting-state and task MRI data in the exploration and replication datasets. Critically, individual differences in intrinsic fluctuations of brain function could lead to spurious FC in group statistics. To conquer this problem, we ran the time-series extraction not only on task EPI, but also on the resting-state EPI of the same participants. By reflecting the individual task-independent FC, correlational maps of resting-state time-series represent a subject-specific baseline for the lag-based analysis. Given the large spatial extent of the network related to the cue phase, different aspects of this phase might be encoded in spatially distinct subregions. Therefore, the extracted time-series of each voxel inside this mask were analyzed separately to allow for regional differences. Building on results of previous studies, the masks related to the picture phase on the other hand were distinctively localized (Walter et al., 2009). Thus, we calculated mean BOLD time-series by averaging the time-series across all voxels inside each of the two masks (pgACC and dmPFC).

### Cross-correlation analysis

Based on the temporal structure of the expectancy paradigm, we used the voxel-based cue-related time-series as seeds in our cross-correlation approach. The picture phase ROIs served as the connectivity targets. To get an estimate of effective or causal connectivity, dynamic causal modeling could be used (DCM, Friston, Harrison, and Penny (2003)). DCM however, assumes causal effects to happen in succession which does not fit with the temporal structure of our task where the difference between cue and picture processing amounts to several seconds. Given the different TR of the exploration and the replication datasets, different lag steps were used for each of the studies to get a maximum time shift of 10 seconds. Imposed by the paradigm, the onset of the picture presentation takes place 3-5 seconds after the cue. Due to the sluggish nature of the BOLD response, we thus expected the corresponding effect within this time window with a peak around 4 seconds. Consequently, the cue-based resting-state and task time-series were cross-correlated with each of the respective picture phase (pgACC, dmPFC) mean time-series using lag steps from −5 to 5 TR for the exploration datasets (5 × 2 TR = 10 seconds) and lag steps of −8 to 8 TR (8 × 1.25 TR = 10 seconds) for the replication datasets. Time-shifted FC values were z-transformed to achieve a compression to a value range between −1 and 1 and normalize the distribution. To ensure that differences in FC arise primarily due to the task demands as described in the previous section, resting-state FC values were subtracted from task FC values yielding delta functional connectivities (dFC, *Figure 1C*). The dFC values were stored as a 3D image at their respective seed voxel coordinates from the cue mask.

## Statistical analysis

### Identification of regions exhibiting a modulatory connectivity profile with the picture processing ROIs

Since we hypothesized to see information transfer from the cue phase to the picture processing phase, we reasoned that regions varying in activation with regard to the respective modality (task or rest), the ROI (pgACC or dmPFC) and lag (time shift in the direction concordant with the paradigm vs time shift in the non-concordant direction) would exhibit a key modulatory role. To test for regions within the cue mask that exhibit this modulatory connectivity profile with the picture phase ROIs in task over rest, we then submitted the 3D maps containing the dFC values to SPM as input for two factorial models (one for each study). Specifically, we tested for regions showing an increased activation for the time shift of 4 seconds that represented the average duration between cue and picture phase versus a time shift of 4 seconds in the negative direction (thus, shifting the cue phase further away from the picture phase). Hence, we set up the models to include the factors modality (task, rest), ROI (pgACC, dmPFC) and lag in seconds (4, −4) and tested for a directed, positive three-way-interaction between modality, ROI and lag.

Given that our dFC maps only contained values within the predefined cue mask, using the standard SPM procedure would have led to incorrect results as thresholding based on random field theory is not applicable to incontinuous masks. AlphaSim was used to simulate the probability of a field of noise based on our cue mask producing a cluster of a given size after the noise has been thresholded at α ≤ .001. We then compared the cluster sizes obtained from the three-way-interaction in SPM to the AlphaSim chance level to identify significant clusters at a corrected α ≤ .05.

Moreover, to test the combined probability of clusters significant in both studies to arise from chance (a conjunction), we used Fisher’s combined probability test. This method combines the p-values from independent test results into one test statistic which follows a X^2^ distribution and thus allows to calculate a conjoint chance probability across both studies.

### Modeling lag-dependent connectivity changes reflecting AIC modulation

As the AIC was the only region showing a preferential modulation of time-shifted connectivity with the two picture processing ROIs (pgACC & dmPFC) in the factorial model, we set up a follow-up analysis to test for differential changes in connectivity profiles over the computed lag steps. In this in-depth analysis of signaling dynamics, we computed the time-shifted connectivity from the AIC (based anatomically on the Hammers atlas; Faillenot, Heckemann, Frot, and Hammers (2017)) to the picture processing ROIs. To investigate if dFC changes over time between AIC and pgACC can be explained by an increase in time shift as would be imposed by the time shift between cue and picture processing phase, we set up a hierarchical linear model where we predicted the time-shifted dFC between AIC and pgACC.

Hierarchical linear models are analogous to full mixed-effects models. They account for the nested nature and corresponding dependency of experimental conditions while allowing for variability in responses both within- and between subjects. For concision, models will be described in terms of their explanatory value for the question at hand (for a more detailed description, see SI). As predictors we included the linear lag steps in seconds (from −10 to 10 seconds) as well as the squared lag steps to model the effect of time shift between cue and picture processing phase. We further added the dFC between AIC and dmPFC on each lag step to control for inter-individual differences in brain responses. Participants were modeled as a random effect to estimate deviations of each individual from the group average. The data hierarchies were then combined using HLM7 (Raudenbush, Bryk, Cheong, Congdon, & du Toit, 2011) for parameter estimation using restricted maximum likelihood.

### Modeling single-trial cue responses as specific predictors of time shifted affective picture processing

By employing cross-correlation analyses on dFC maps, we investigated if a time lag congruent with the paradigms’ temporal structure is evident in the connectivity profile of SN nodes with picture processing regions. Given that the entire time-series is used for the cross-correlation analysis, this, however, is not yet sufficient to establish the critical role of cue processing. Hence, building on these previous results, we wanted to assess the stronger hypothesis that time shifted changes in connectivity are specifically following cue phases in the paradigm with the next set of analyses. If the AIC acts as the hypothesized switch, it should exert a modulatory influence on the task-positive and task-negative nodes preferentially for the trials comprising a cue as these deliver the predictive information necessary for switching. Moreover, the magnitude of the cue response in the AIC should be predictive of the lagged activation in the target regions over and above the activation in the target brain region (i.e., due to autocorrelation of the signals). To test this stronger version of the hypothesis, we first needed to estimate single-trial level brain responses employing a second set of HLM analyses (Kroemer et al., 2014; Kroemer et al., 2016). For this first model, we extracted the regressors for the cue phase, the presentation of an expected picture, and the presentation of an unexpected picture from the unfiltered design matrix of the SPM first-level statistics. Based on these regressors, we then predicted the AIC time series to estimate single-trial beta weights of AIC signal for all trials containing a cue as we hypothesized the AIC to act as a switch specifically in response to the cue signal.

Next, we set up the second part of this HLM analysis to test if a lagged connectivity modulation from the AIC to the dmPFC preferentially occurs for AIC signal increases during the cue phase which would indicate that the AIC based switching is not driven by general task demands. The dmPFC is associated with task-positive processing and is considered to be the target for the cue related information forwarded by the AIC to enhance picture processing 4 seconds later. Thus, we set up the single-trial AIC cue-response weights estimated in the previous model to predict the dmPFC signal lagged by 4 seconds. Critically, the unshifted time series of the dmPFC was also added to control for autocorrelation within the time series. Hence, a predictive effect of the estimated single-trial cue responses on the time shifted dmPFC signal in this case would indicate a transfer of information over time rather than a mere effect of information redundancy in the BOLD responses as variance due to autocorrelative effects is already removed.

## Results

### AIC channels task-specific functional connectivity

To elucidate how top-down expectancy signals modulate future affect, we analyzed two independent samples where participants viewed cued and uncued emotional pictures. First, we sought to identify regions within the cue mask (see SI) that exhibit a modulatory connectivity profile with the prefrontal ROIs involved in affective processing. A factorial model showed a significant Modality (task, rest) × ROI (pgACC, dmPFC) × Lag (4, −4) interaction in the AIC (*Figure 2*; EXP: T = 3.66, *p*_corrected_ = 0.047; REP: T = 4.84, *p*_corrected_ = 0.035, conjunction of whole-brain corrected p-values: *p*_EXP∩REP_ = 0.012). Thus, the AIC was the only region modulating the two picture processing ROIs lagged task-specific functional connectivity in opposite directions.

**Figure 2.**
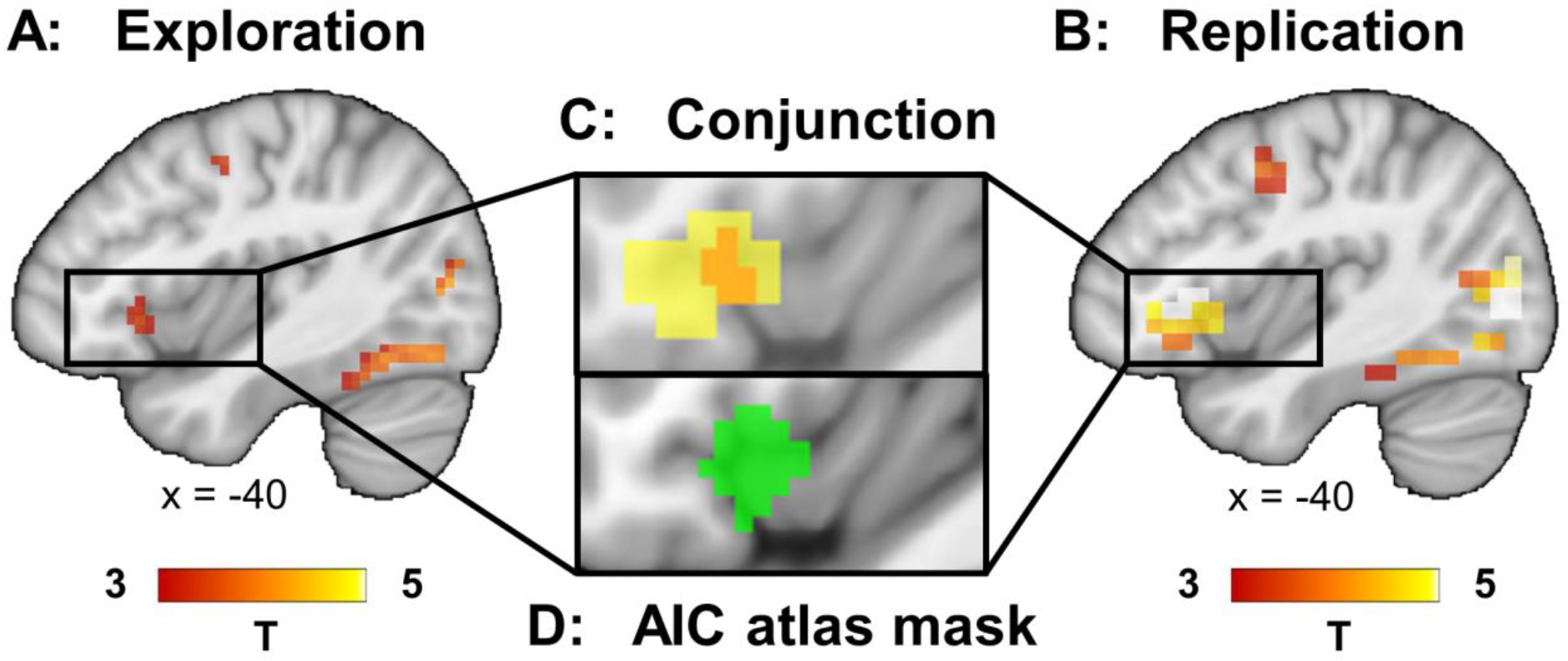
Results from a factorial second-level model on the individual functional connectivity maps, showing that the anterior insula differentially modulates task-specific functional connectivity with the two picture processing ROIs (pgACC and dmPFC). The model included the factors Modality (task, rest), Region of interest (pgACC, dmPFC) and Lag in seconds (4, −4). Depicted here are thresholded T-maps (α ≤ 0.001, uncorrected) of the Modality × Region × Lag interaction in the anterior insula for (A) the Exploration study, (B) the Replication study and (C) the overlap between both studies’ maps. For comparison, (D) depicts the anterior insula (AIC) mask based on the Hammers atlas (Faillenot et al., 2017) used in the follow-up analyses.

### AIC modulates lag-dependent FC differentially for task-positive and task-negative nodes

To investigate the temporal modulation of task-specific connectivity between AIC and the picture processing ROIs, hierarchical linear models were estimated (*Figure 3*). Thus, we predicted the time-lagged delta connectivity between the AIC and the pgACC using linear and squared lag as predictors. Furthermore, we added the time-lagged delta connectivity between the AIC and the dmPFC to control for inter-individual differences in connectivity within this network. This revealed a significant effect of the linear lag (*Table 1*; EXP: T_df=36_= −2.787, *p* = 0.008; REP: T_df=54_= −2.7, *p* = 0.009, conjunction *p*_EXP∩REP_< 0.001) and AIC – dmPFC dFC (EXP: T_df=36_= 5.821, *p* < 0.001; REP: T_df=54_= 10.047, *p* < 0.001, conjunction *p*_EXP∩REP_ < 0.001). The squared lag component was only significant in the exploration study (EXP: T_df=36_= 3.299, *p* = 0.002; REP: *p* = 0.112). Thus, in line with the temporal structure imposed by our paradigm, task-specific functional connectivity between the AIC and picture processing nodes is modulated over several steps of time shift.

**Table 2.** Results from Hierarchical Linear Models predicting delta functional connectivity (task-rest) between the pgACC and the AIC for (A) the Exploration study and (B) the Replication study show, that time shift predicts changes in functional connectivity profiles.

|  | Coefficient | T | p |
| --- | --- | --- | --- |
| Intercept | -0.057 | -3.517 | 0.001 |
| Lag linear | -0.004 | -2.787 | 0.008 |
| Lag squared | 0.001 | 3.299 | 0.002 |
| dmPFC <> AIC dFC | 0.468 | 5.821 | < 0.001 |
| df = 36 |  |  |  |

**(B)** Replication study
|  | Coefficient | T | p |
| --- | --- | --- | --- |
| Intercept | -0.016 | -1.620 | 0.111 |
| Lag linear | -0.012 | -2.700 | 0.009 |
| Lag squared | 0.025 | 3.506 | < 0.001 |
| dmPFC <> AIC dFC | 0.554 | 10.047 | < 0.001 |
| df = 54 |  |  |  |

**Note.** Models included an intercept and the linear and squared lag in seconds as predictors while the delta functional connectivity (task-rest) between the dmPFC and the AIC was added to control for inter-individual differences in brain response.

**Figure 3.**
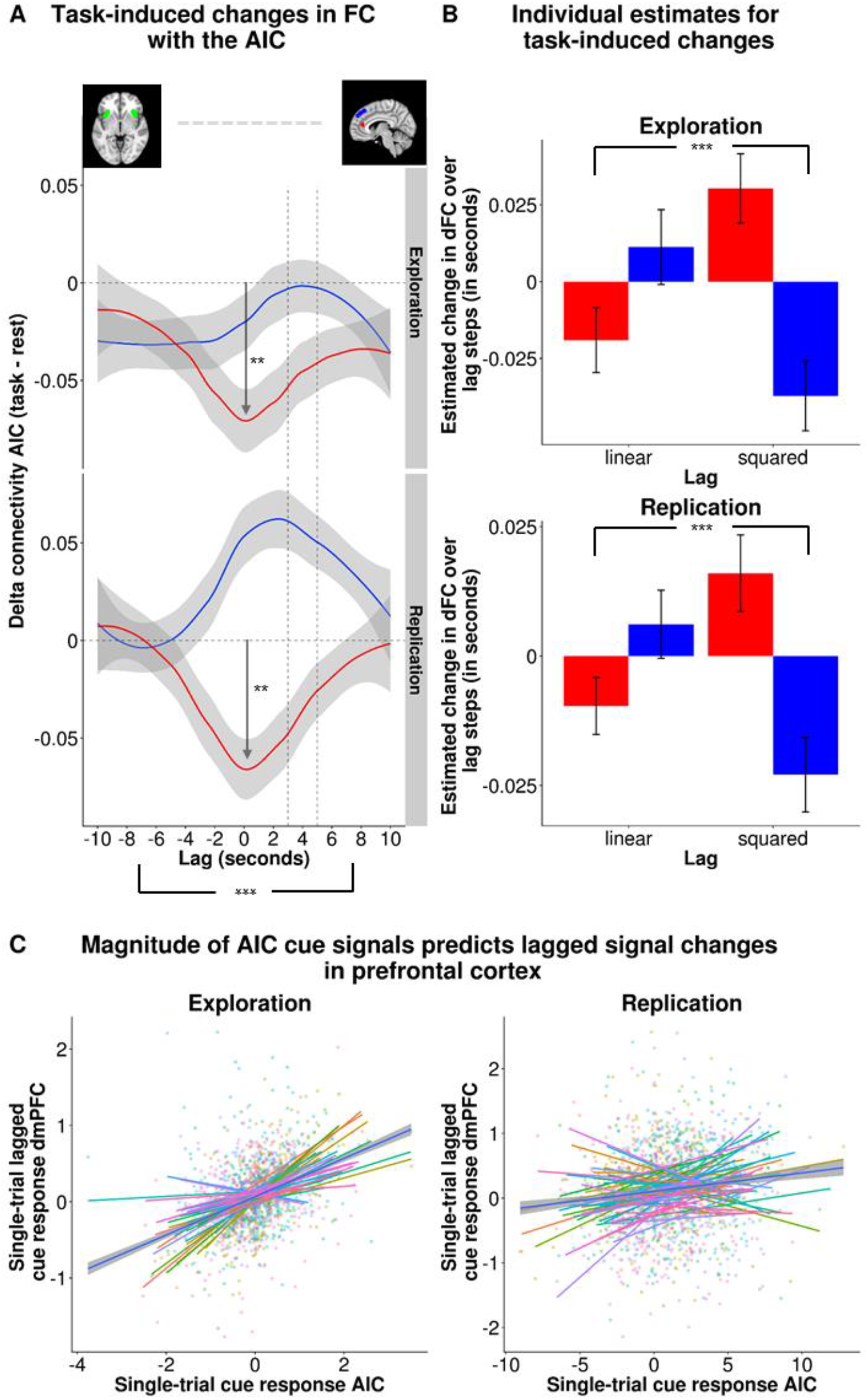
(A) Delta functional connectivity (task-rest) between the AIC and the two picture processing ROIs pgACC (red) and dmPFC (blue) over the lag steps of −10 to 10 seconds in the Exploration study (upper panel) and the Replication study (lower panel). Vertical striped lines indicate the lag window of 3-5 seconds that corresponds to the delay between cue presentation and onset of picture presentation. (B) Significant interaction between lag (linear and squared) and estimated change in delta functional connectivity (task-rest) between the AIC and pgACC (red) as well as dmPFC (blue) shows that delta functional connectivity for the two picture processing ROIs is modulated in opposite directions for both studies. While delta functional connectivity between the AIC and pgACC decreases before it returns to baseline, delta functional connectivity between the AIC and dmPFC increases in anticipation of task demands. (C) Single-trial estimates of the AIC cue response and the lagged dmPFC cue response including the unshifted dmPFC cue response and uncued trials as a covariate, show that the AIC based switching is specific for expected picture processing. A higher response of the AIC to the cue is associated with a higher response of the dmPFC to the cue after it has been shifted by 4 seconds which corresponds to the delay between cue presentation and onset of picture presentation.

### Lag-dependent FC changes are associated with signals of the AIC specifically during cue phase

After we showed a replicable link between time-lagged task-specific functional connectivity in line with the temporal prediction of cue processing, we tested more specifically if the magnitude of the cue response in the AIC is predictive of the time-shifted activation in the dmPFC over and above the unshifted dmPFC activation. We found that single-trial betas for cue signals in the AIC significantly predicted time-shifted dmPFC activation (*Table 2*, EXP: T_df=36_ = −5.704, *p* < 0.001; REP: T_df=54_ = −6.269, *p* < 0.001, conjunction *p*_EXP∩REP_ < 0.001). Hence, the results of this single-trial model corroborate the notion that the AIC based switching and modulation of task-positive and task-negative network nodes is restricted to AIC signal increases in response to cues (*Figure 4 & 5*).

**Table 2.** Results from Hierarchical Linear Models predicting dmPFC BOLD signal lagged by 4 seconds for (A) the Exploration study and (B) the Replication study show, that the single-trial AIC response to cues predicts the lagged dmPFC signal beyond mere effects of autocorrelation.

|  | **Coefficient** | **T** | **p** |
| --- | --- | --- | --- |
| Intercept | 0.007 | 0.617 | 0.541 |
| AIC cue beta | -0.122 | -5.704 | < 0.001 |
| Unshifted dmPFC time-series | 0.210 | 8.942 | < 0.001 |
| df = 36 |  |  |  |

**(B)** Replication study
|  | **Coefficient** | **T** | **p** |
| --- | --- | --- | --- |
| Intercept | -0.018 | -1.625 | 0.110 |
| AIC cue beta | -0.095 | -6.269 | < 0.001 |
| Unshifted dmPFC time-series | 0.207 | 11.004 | < 0.001 |
| df = 54 |  |  |  |

**Note.** Models included an intercept and the beta weight for AIC signal during cue phases as predictors while the unshifted dmPFC signal was added to control for autocorrelation.

**Figure 4.**
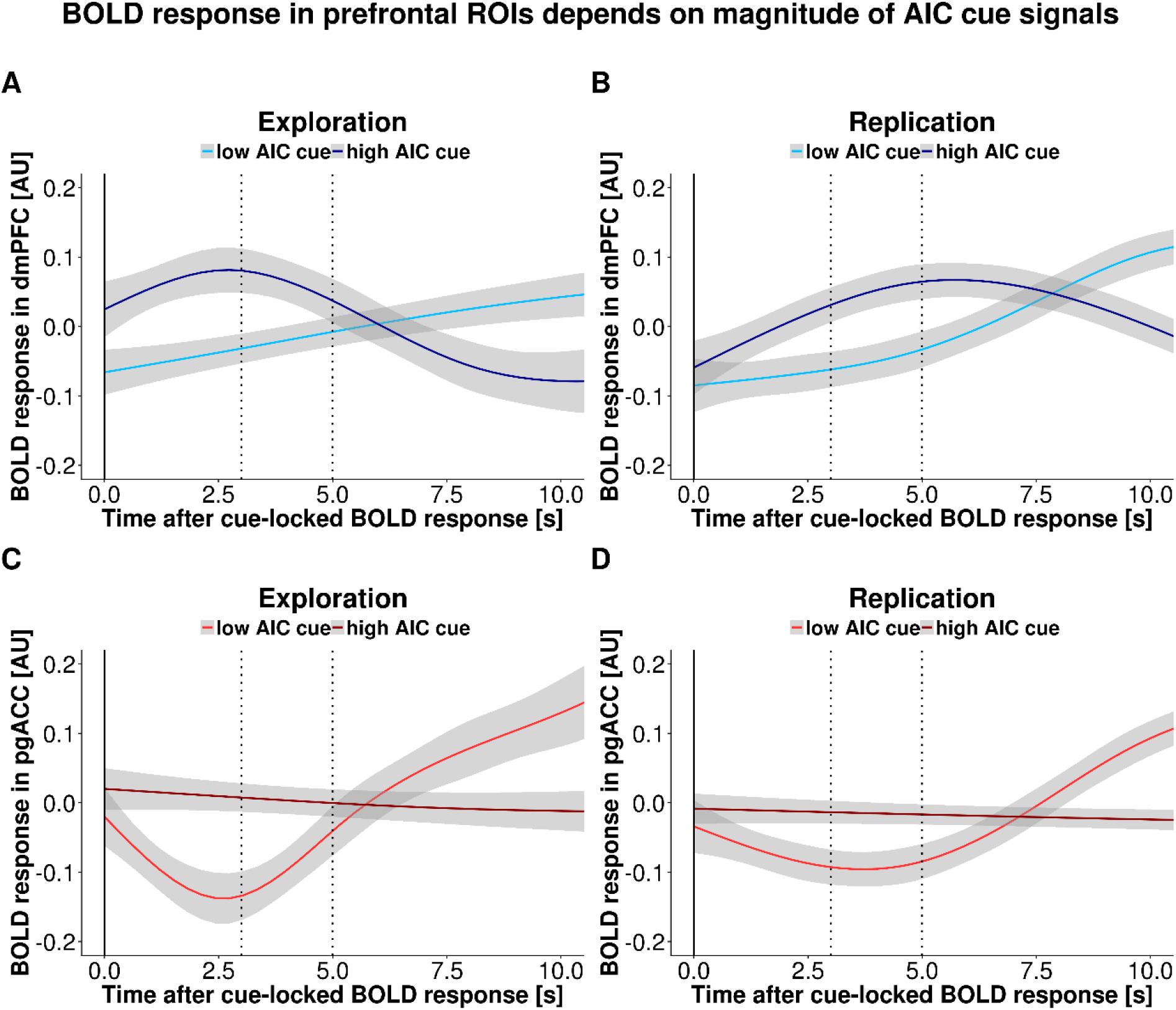
BOLD response of the two picture processing ROIs dmPFC (A & B) and pgACC (C & D) for the Exploration (A & C) and Replication (B & D) study depending on either a high or low BOLD signal of the AIC during the cue phase. On the x-axis, the time in seconds after cue onset is plotted. Vertical striped lines indicate the lag window of 3-5 seconds that corresponds to the delay between cue presentation and onset of picture presentation. While a high AIC signal is associated with an increase in the dmPFC signal in the target lag window, a low AIC signal leads to a substantially delayed signal increase in this task-positive region. Contrary to that, the pgACC signal is modulated specifically for the low AIC signal.

**Figure 5.**
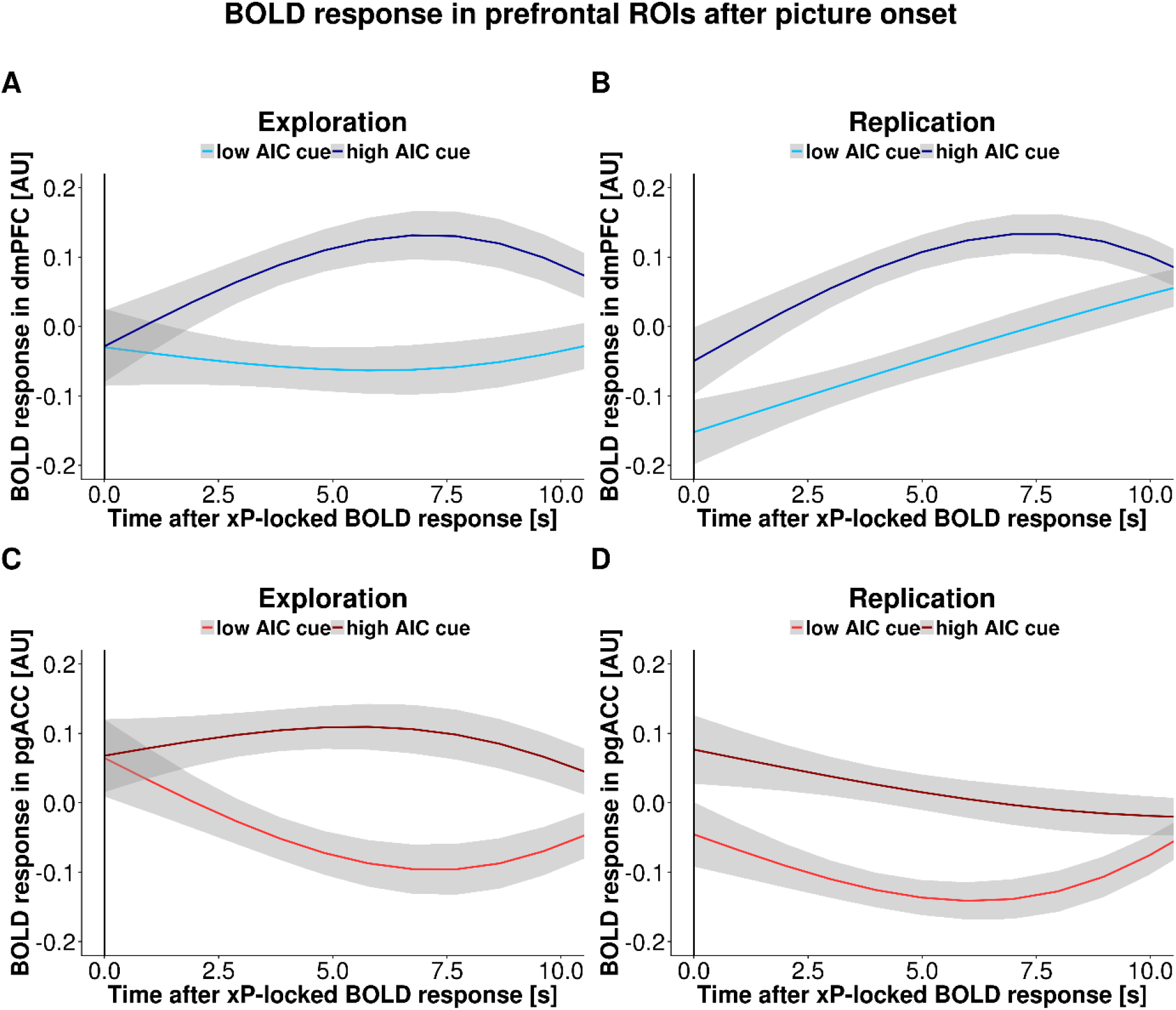
BOLD response of the two picture processing ROIs dmPFC (A & B) and pgACC (C & D) for the Exploration (A & C) and Replication (B & D) study depending on either a high or low BOLD signal of the AIC during the cue phase. On the x-axis, the time in seconds after the onset of a cued picture is plotted. A high AIC cue signal is associated with an increase in both dmPFC and pgACC signal suggesting that cue information is preserved into the picture phase given the cue was processed actively within the AIC.

## Discussion

Expectations modulate impending affective responses, yet it was unknown how this tuning is implemented and transferred from the processing of cues to emotional pictures in the brain. Using cross-correlation analysis of BOLD time series, we found that expectancy signals in the AIC channel salience, leading to differential coupling with prefrontal networks related to task-positive (dmPFC) and task-negative (pgACC) processing. Critically, we observed that stronger cue signals in the AIC predicted lagged signal increases in the dmPFC (controlled for its autocorrelation) indicating neural traces of expectations elicited by the cue. Moreover, these temporal profiles of AIC-based modulation were robust across two independent studies cueing the impending presentation of affective pictures. Thus, our results shed new light on the putative role of the AIC as a hub of the SN implementing dynamic switching between prefrontal networks implementing task-positive and task-negative processing.

The AIC has been previously hypothesized to gate information processing and our results corroborate this interpretation by demonstrating that the AIC channels the recruitment of differential prefrontal networks depending on salience elicited by the cue. This is also well in line with the functional specialization and resulting network interactions described in the EPIC model (Feldman Barrett & Simmons, 2015). Feldman Barrett and Simmons (2015) argue that agranular regions such as the AIC and the pgACC primarily generate predictions while granular regions such as the dmPFC compare received sensory signals with predictions to compute prediction errors. Thus, our results corroborate the idea that cytoarchitecture can inform computational principles within extended neural networks such as expectancy-based channeling of affective processing according to principles of active inference. Notably, we showed that the synchronization between regions generating (AIC, pgACC) versus updating (dmPFC) predictions is lagged in correspondence with the interval between phases of cue and picture presentation. Collectively, this suggests that the lagged connectivity may reflect traces of active inference where the AIC generates predictions that channel future processing of affective pictures.

Critically, our results also show that the lagged functional connectivity between AIC and prefrontal networks did not result from the fixed temporal structure of the paradigm as lagged signal changes in the dmPFC were predicted from the magnitude of the cue-induced signal in the AIC. This result reemphasizes influential ideas that the cue-induced response magnitude of the AIC reflects the expected saliency of incoming stimuli. Such saliency encoding may directly serve the channeling (Dehaene, Changeux, Naccache, Sackur, & Sergent, 2006; Dehaene & Naccache, 2001) or “gating” function (Michel, 2017) assigned to the AIC determining how sensory input as information is forwarded for further processing to higher-order prefrontal areas. Likewise, the global neuronal workspace hypothesis (Dehaene & Naccache, 2001) posits that top-down influence needs to be exerted for information to become conscious and represented within a network of fronto-parietal regions with long-distance connections (global neuronal workspace). Moreover, the central role of the AIC across a wide range of tasks and functions (Uddin, Nomi, Hebert-Seropian, Ghaziri, & Boucher, 2017), its stable hub-like network properties (Gollo et al., 2018; van den Heuvel & Sporns, 2011), and the converging inputs from different sensory modalities (Butti & Hof, 2010; Downar, Crawley, Mikulis, & Davis, 2000; Evrard, Logothetis, & Craig, 2014) make the AIC a prime candidate for channeling expectations. Such rapid long-distance integration of cortical afferents may be supported by strongly localized *von Economo* neurons that have been identified in the primate AIC (Evrard, 2018) and might enable the synchronization between networks such as SN, CEN, and DMN. Intriguingly, the function of the AIC and ACC is also most commonly affected across neuropsychiatric disorders (Goodkind et al., 2015), particularly in major depressive disorder (DeVille et al., 2018; Iwabuchi et al., 2014; Namkung, Kim, & Sawa, 2017) pointing to a vital role of keeping expectations and outcomes in sync (Feldman Barrett & Simmons, 2015). Thus, our results support the notion that the magnitude of the cue-induced signal in the AIC reflects a salience estimate necessary to subsequently engage higher-order processing areas like the dmPFC.

Furthermore, our results show that by channeling cue-related information the AIC also induces a switch between DMN and CEN network nodes. While a wealth of studies has been conducted investigating the effects of expectation on subsequent (affective) processing using regressor based analyses of the separated stages of the paradigm (Bermpohl et al., 2006a, 2006b; Bermpohl et al., 2009; Blair et al., 2007; de Lange et al., 2018; Northoff et al., 2004; Pessoa, 2008, 2016; Walter et al., 2009), our proposed method of using cross-correlations on fMRI time-series allows to track the information transfer between key functional networks over time by exploiting the timing information imposed by our task. Interestingly, our results suggest, that an increase in the AIC signal during the cue phase modulates the signal in the dmPFC, while the pgACC is modulated specifically for low AIC cue-signals. Expanding the view on the commonly reported network interplay in resting-state (Bressler & Menon, 2010; Menon, 2011; Seeley et al., 2007; Uddin, 2015), this indicates that the SN and DMN may be synchronized by default. This notion is also supported by studies showing a combined network of the AIC and ACC inducing switching between the DMN and CEN (Sridharan, Levitin, & Menon, 2008) as well as improved performance for attention switching based on increased DMN activity before task execution (Bogler, Vowinkel, Zhutovsky, & Haynes, 2017). Hence, a low signal of the AIC may be a precondition for the DMN to be active as the basic state the brain reverts to in absence of any external task demands until an increased AIC signal induces the switching from an active DMN to an active CEN in light of emerging task demands.

Nevertheless, our results are limited by several factors which need to be addressed in future studies. First, as we were interested in the general mechanism of expectancy channeling, we did not differentiate between the valence categories of stimuli. Recent studies modeling mood fluctuations indicate AIC modulation might partially depend on stimulus valence (Vinckier, Rigoux, Oudiette, & Pessiglione, 2018). However, replication of our results in the second dataset containing no negatively-valenced stimuli suggests that the lagged channeling of prefrontal networks via the AIC can be seen as a generalizable mechanism in cued processing. Second, the characterization of the AIC as one homogeneous agranular region generating prediction as proposed in the EPIC framework might be too simplistic (Evrard et al., 2014). However, limitations in spatial and temporal resolution with current imaging sequences impede the functional localization of such fine-grained details in humans to date. Thus, high-resolution laminar fMRI techniques may allow to shed light on layer-specific origins of prediction and prediction-error signals to further delineate expectancy-based channeling of sensory input.

To conclude, expectancy shapes our perception of impending events and aberrant expectations are well-known to play an important role in affective disorders. Based on cross-correlation analysis and single-trial hierarchical linear modeling of BOLD responses, we have shown that cue signals in the AIC channel salience, implementing a switch in functional coupling with either the DMN (as indexed by the pgACC) or the CEN (as indexed by dmPFC). Collectively, our results suggest that the SN and DMN are synchronized “by default”. As the AIC monitors and integrates external information to determine the priority of external demands, it orchestrates the switch of prefrontal functional coupling from internally-oriented to externally-oriented processing. Thereby, our results add a crucial novel facet to the understanding of signaling dynamics supporting the interplay of top-down expectations and bottom-up affective input by identifying the AIC as a critical switch channeling the subsequent affective reception of events.

## Acknowledgements

VT & NBK received salary support from the University of Tübingen, Faculty of Medicine fortüne grant #2453-0-0. MPN & NBK received salary support from the Else Kröner-Fresenius-Stiftung, grant #2017-A67. MW was supported by DFG grant (Wa2674/4-10) and SFB779-A06. The Exploration dataset was collected along with EEG data in a clinical trial sponsored by Biologische Heilmittel HEEL GmbH, Germany (NCT02602275) in which MW was a PI. fMRI data was provided as courtesy for the purpose of these analyses which were not related to the trial objectives. The authors thank Shijia Li for her contribution regarding data acquisition and prior analyses within the Replication study. The authors declare no conflict of interest.

## Author contributions

MW was responsible for the concept and design of the studies. JVDM, VB & LH collected data. NBK & VT conceived the method and VT & YF processed the data. VT performed the data analysis and NBK & MW contributed to analyses. VT & NBK wrote the manuscript. All authors contributed to the interpretation of findings, provided critical revision of the manuscript for important intellectual content and approved the final version for publication.

## Data and Code availability statement

The codes and full data of the study are currently not publicly available due to ongoing additional analyses, however, individual summary data and codes concerning this manuscript may be available on reasonable request from the authors.

