## Supplementary Information for "The anterior insula channels prefrontal expectancy signals during affective processing"

Vanessa Teckentrup[1], Johan N. van der Meer[2-5], Viola Borchardt[4,5], Yan Fan[6], Monja P. Neuser[1], Claus Tempelmann[7], Luisa Herrmann[1], Martin Walter[1,3-5*#], Nils B. Kroemer[1*#]

[1] University of Tübingen, Department of Psychiatry and Psychotherapy, Tübingen, Germany

[2] Queensland Institute of Medical Research, Brisbane, Australia

[3] University of Magdeburg, Department of Psychiatry and Psychotherapy, Germany

[4] Clinical Affective Neuroimaging Laboratory, Magdeburg, Germany

[5] Leibniz Institute for Neurobiology, Magdeburg, Germany

[6] Leibniz Research Centre for Working Environment and Human Factors, Department of Psychology and Neurosciences Dortmund, German

[7] University of Magdeburg, Department of Neurology, Germany

### contributed equally

Corresponding authors*:

Prof. Dr. Martin Walter | Dr. Nils B. Kroemer

Department of Psychiatry and Psychotherapy

University of Tübingen

Calwerstr. 14

72076 Tübingen, Germany

#### Participants and procedure

Study 1 (EXP) was conducted as a randomized, placebo-controlled, double blind crossover clinical trial to test for effects of Neurexan (Nx4), a medicinal product, consisting of three herbal extracts (Avena sativa, Coffea arabica, Passiflora incarnate) and one mineral salt (Zincum isovalericum). Each participant was invited to the lab on two days. Each measurement started with a simultaneous EEG-fMRI session in which anatomical (5 minutes) and baseline resting-state measurements (R1, 12 minutes) were conducted. Then participants received either verum or placebo depending on the day and their randomization scheme. This was followed by an EEG only session where participants underwent two tasks assessing attention (attentional modulation of salience task & oddball task). After this, the second resting-state simultaneous EEG-fMRI measurement was conducted (R2, 12 minutes) followed by simultaneous EEG-fMRI measurements of the Hariri task (8 minutes), the emotional expectancy task (15 minutes) employed for the analyses described in this paper, the Scan Stress task (15 minutes) and a third resting-state measurement (R3, 12 minutes) at the end.

In Study 2 (replication: REP), 86 participants (24 female) were investigated. The study comprised the acquisition of patients suffering from major depressive disorder (n = 30) as well as healthy control participants (n = 56). For the analyses reported in this paper, only data from healthy controls was used. Again, one participant was not included in the final sample due to excessive head motion leading to the final sample size of n = 55 (13 female) with a mean age of 32.1 (± 8.5, range: 22-52) for the replication study. Participants in Study two first underwent an anatomical MRI scan (10 minutes) followed by 5 magnetic resonance spectroscopy scans (9 minutes each). Afterwards, participants completed a resting-state fMRI measurement (10 minutes) and the salience expectancy measurement (14 minutes) both of which were used for the current analyses.

#### Paradigm

##### Replication study

In our replication study we employed a paradigm designed to discern expectancy effects related to high and low salient picture stimuli. The factorial design comprised the two factors *expectancy* (expected, unexpected) and *salience* (high, low). Every trial started with a fixation cross which was shown for a varying duration of 7.5-10.5 seconds. In half of the trials, the fixation cross was followed by a cue phase, in which the salience of the subsequently presented picture was cued by either an upwards pointing arrow with an exclamation mark (high salience) or a downwards pointing arrow with an exclamation mark (low salience). The cues were shown in white on a black background and for a varying duration of 3-5 seconds. In the subsequent picture phase, a picture was shown in accordance to the salience indicated by the cue. In half of the trials, unexpected pictures were not preceded by a cue phase but were shown directly after the fixation cross. In total, 40 pictures from the International Affective Picture System (Lang, Bradley, & Cuthbert, 2008) were used with positive (high salience) and neutral (low salience) pictures shown for 4 seconds in a counterbalanced design (i.e., 20 pictures per salience category). The paradigm also comprised cues that did not show an arrow but either one or two dots cueing whether one or two persons will be depicted on the subsequent picture. For the analyses reported in this paper however, only the salience related cues and subsequent pictures were of interest and therefore analyzed. Completing the paradigm took about 14 minutes.

#### MRI data acquisition and preprocessing

##### MRI data preprocessing

For the exploration study, fMRI task data was submitted to SPM12 (Statistical parametric mapping, Wellcome Department of Imaging Neuroscience, London, UK; http://www.fil.ion.ucl.ac.uk/spm/software/spm12/) implemented in Matlab R2017a (The Mathworks Inc., Natick, MA, USA). First, slice timing was corrected for each volume by interpolating the slices to the middle slice, then all volumes were realigned to the first volume by applying a rigid-body transformation to correct for head motion. Participants with head movement exceeding 3 mm for translation head motion parameters or 3° for rotation head motion parameters were excluded from further analyses. The anatomical images were coregistered to match the functional images. All images were then spatially normalized to standard Montreal Neurological Institute (MNI) space following tissue segmentation of the anatomical images and subsequent application of the generated deformation field to the functional images. All images were smoothed with a Gaussian kernel with 8 mm FWHM.

For the replication study, fMRI task data was submitted to SPM12 implemented in MATLAB R2017a. The first 8 volumes were discarded to allow the MR signal to achieve T1 equilibrium. First, slice timing was corrected for each volume by interpolating the slices to the middle slice, then all volumes were realigned to the first volume by applying a rigid-body transformation to correct for head motion. Participants with head movement exceeding 3 mm for translation head motion parameters or 3° for rotation head motion parameters were excluded from further analyses. The anatomical images were coregistered to match the functional images. All images were then spatially normalized to standard Montreal Neurological Institute (MNI) space following tissue segmentation of the anatomical images and subsequent application of the generated deformation field to the functional images. All images were smoothed with a Gaussian kernel with 6 mm FWHM.

Resting-state fMRI data within both studies was preprocessed using MATLAB R2017a, SPM12 and DPABI v2.1 (Data Processing & Analysis for Brain Imaging toolbox; http://rfmri.org/dpabi). The first 5 volumes were discarded to allow the MR signal to achieve T1 equilibrium. Then, slice timing was corrected for each volume by interpolating the slices to the middle slice and all volumes were realigned to the first volume by applying a rigid-body transformation to correct for head motion. The anatomical images were coregistered to match the functional images, then segmented into gray matter (GM) and white matter (WM). Subject-specific templates were created with diffeomorphic anatomical registration using DARTEL (Ashburner, 2007). A group-specific template was then created from all subject-specific templates. Coregistered rs-fMRI data were subsequently normalized to the MNI template. Physiological noise was reduced by regressing out signals from white matter (WM), cerebrospinal fluid (CSF) and the 6-rigid body realignment parameters. All images were smoothed with a Gaussian kernel with 8 mm FWHM for the exploration study or 6 mm FWHM for the replication study, respectively. Importantly, global signal removal was not performed to avoid false induction of anti-correlations between time-series (Murphy, Birn, Handwerker, Jones, & Bandettini, 2009).

#### Definition of regions of interest (ROI)

##### *Cue mask*

To capture the representation of general expectancy (collapsed over valence) during the cue phase, a mask was defined that represented voxels showing activation above the significance threshold of $T = 3.11$ ($p < 0.001$, uncorrected) within the cue phase. First, the preprocessed datasets from the exploration study were submitted to SPM12. Onsets and durations for each cue phase were entered to subsequently convolve these design regressors with the canonical HRF leading to the output of task-based regressors. To avoid overfitting the masks to the individual expectancy contrast, T-maps were calculated for each subject for the contrast cue > baseline in a leave-one-out cross-validation approach. Hence, beta maps of this contrast of interest of all participants minus one were averaged to create the map for the left- out participant. Subsequently the T-map was calculated by dividing this mean beta map by the standard error of all included beta maps. The T-map was then thresholded according to two criteria: Clusters where only included if they exceeded a size of 50 voxels with a voxel-based threshold of $T = 3.11$ (i.e., $p < 0.001$, uncorrected). After completion of this leave-one-out process, a group mask was generated by selecting only those voxels that were present in each thresholded T-map over all participants. This yielded a mask comprising a volume of 139.95 $cm^3$ (5170 voxels) for the exploration study. For the replication study, we resliced this mask using the SPM12 reslice function with nearest neighbor interpolation which yielded a mask comprising a volume of 143.25 $cm^3$ (1146 voxels).

***pgACC***

To investigate how the magnitude of the cue representation influences picture processing, we extracted the median beta value from the voxels within the cue mask (defined independently via the leave-one-out cross-validation). To uncover brain areas where activation during the processing of expected pictures covaried with the magnitude of the signal during the cue presentation, we then used this proxy of the magnitude of cue representation in a subsequent linear regression model. Then, we calculated the linear association of cue representation with the contrast expected picture viewing > unexpected picture viewing. Among other regions, this yielded a cluster in the pregenual anterior cingulate cortex (pgACC) that was characterized by a negative correlation between the elicited magnitude of the cue signal and expected picture processing. As this cluster represented the task-negative aspect of the picture processing and the pgACC is a prominent node of the default mode network (Uddin, 2015), we subsequently exported this cluster as a ROI mask for the picture phase to represent the internally-driven aspects in our analysis.

***dmPFC***

We hypothesized that the dmPFC plays a key role in handling affective picture processing given its continuous recruitment in simultaneous processing of cognitive and emotional demands (Bermpohl et al., 2006; Walter et al., 2009). Based on this, we selected a second ROI in the dorsomedial prefrontal cortex (dmPFC). This ROI was based on Walter et al. (2009), who investigated the functional subdivisions of the dmPFC in processing expected stimuli using emotionally as well as erotically salient pictures in an expectancy task. Following the procedure detailed by Walter et al. (2009), we drew 6 spheres with a diameter of 5 mm using the WFU pick atlas (https://www.nitrc.org/projects/wfu_pickatlas/) implemented in MATLAB 2016a (The MathWorks, Natick). These spheres were located bilaterally 5 mm from the midline at the y and z MNI coordinates of [38, 50], [45, 45] and [50, 38]. For the analysis, these spheres were handled as one region of interest labeled as dmPFC. To ensure comparable analysis pathways for the exploration and the replication datasets, all masks were resliced to the dimensions and voxel size of the replication datasets using the SPM12 reslice function with nearest neighbor interpolation.

#### Statistical analysis

##### Modeling lag-dependent connectivity changes reflecting AIC modulation

To set up the lowest level of the model, we incorporated paradigm-related information. We included the linear lag steps in seconds (from -10 to 10 seconds) as well as the squared lag steps to investigate if dFC changes over time between AIC and pgACC can be explained by an increase in time shift as would be imposed by the time shift between cue and picture processing phase. We further added the dFC between AIC and dmPFC on each lag step to control for inter-individual differences in brain responses. The second level of the hierarchy then contained participant level information, modeled as a random effect to estimate deviations of each individual from the group average. The two data hierarchies were then combined using HLM7 (Raudenbush, Bryk, Cheong, Congdon, & du Toit, 2011) for parameter estimation using restricted maximum likelihood.

##### Modeling single-trial cue responses as specific predictors of time shifted affective picture processing

The first model was used to estimate single-trial beta weights of AIC signal for all trials containing a cue. For this, we created the lowest level of the hierarchy by extracting the regressors for the cue phase, the presentation of an expected picture and the presentation of an unexpected picture from the unfiltered design matrix of the SPM first-level statistics. On the second level we allowed single trials to vary, modeling them as a random effect as the brain response to a trial might differ over the course of the paradigm. Analogous to this, we also modeled participant-related differences to vary freely on the third level. The data hierarchies were combined using HLM 7 and full maximum likelihood estimation to predict the extracted task time series of the AIC.

For the second model we also extracted the task time series of the dmPFC and computed the dmPFC signal time-shifted by 4 seconds. For the lowest level of this hierarchy, we included information on the paradigm level, such as the regressor for the cue phase (also used in the prior model) to control for the difference between expected and unexpected pictures. The unshifted time series of the dmPFC was also added to the model to control for autocorrelation within the time series. On the second level of the hierarchy, we included trial-based information and, hence, added the single-trial AIC cue responses estimated in the preceding model. In order to allow the estimated cue signal to vary over trials depending on the processing of the cue we modeled them as a random effect. Lastly, we also included participants as a random effects variable on the third level. Again, we combined the data hierarchies using HLM 7 and full maximum likelihood estimation.
